# Spontaneous Variations in Arousal Modulate Subsequent Visual Processing and Local Field Potential Dynamics in the Ferret during Quiet Wakefulness

**DOI:** 10.1101/2022.05.31.494160

**Authors:** Lianne M.F. Klaver, Lotte P. Brinkhof, Tom Sikkens, Lorena Casado-Román, Alex G. Williams, Laura van Mourik-Donga, Jorge F. Mejías, Cyriel M.A. Pennartz, Conrado A. Bosman

## Abstract

Behavioral states affect neuronal responses throughout the cortex and influence visual processing. Quiet wakefulness (QW) is a behavioral state during which subjects are quiescent, but awake and connected to the environment. Here, we examined the effects of pre-stimulus arousal variability on post-stimulus neural activity in primary visual cortex (V1) and posterior parietal cortex (PPc) in awake ferrets, using the pupil diameter as an indicator of arousal. We observed that during low arousal, low-frequency power increases during visual stimulation, and that the peak alpha frequency shifted depending on the arousal state. High arousal increased gamma power as well as low-frequency inter- and intra-areal coherence. Using a simplified model of laminar circuits, we show that this connectivity pattern is compatible with feedback signals targeting infragranular layers in area PPc and supragranular layers in V1. Neurons in V1 displayed higher firing rates at their preferred orientations on high-arousal trials. Broad-spiking cells in V1 entrained to high-frequency oscillations (>80 Hz), whereas narrow-spiking neurons phase-locked to low (12-18 Hz) and high-frequency (>80 Hz) rhythms. These results indicate that the variability and sensitivity of post-stimulus cortical responses and coherence depend on the pre-stimulus behavioral state and account for the neuronal response variability observed during repeated stimulation.

---

When cortical neurons are repeatedly stimulated, their responses are highly variable (Vogels et al. 1989; Arieli et al. 1996). This variability results from the spontaneous activity of anatomically interconnected neurons throughout the cortex and is associated with the animal’s behavioral state (Ecker et al. 2010; Harris and Thiele 2011; Schölvinck et al. 2015; Denfield et al. 2018; Jacobs et al. 2020). Behavioral states can exert global influences throughout the cortex and distinctively affect neuronal responses and stimulus perceptibility (Montijn et al. 2015). Therefore, understanding the relationship between behavioral states and cortical activity is critical to elucidate how neuronal variability affects cortical processing.

Behavioral states range from highly active and attentive to deep sleep stages. These states correlate with well-defined electrophysiological patterns of brain activity. For example, high-frequency oscillations and desynchronized waves are predominant during wakefulness, whereas during deep sleep, highly synchronized low-frequency activity patterns dominate brain activity (Steriade et al. 1993; Steriade 2006; Harris and Thiele 2011; McGinley, Vinck, et al. 2015; Sanchez-Vives et al. 2017). During wakefulness, animals can transition between periods of attentive activity and wakeful quiescence. These periods of quiet wakefulness (QW) share features with both active wakefulness (e.g., active behavior, locomotion, increased muscular tone) and sleep (e.g., immobility, hippocampal sharp-wave ripples), and their occurrence depends on several factors such as arousal, attention, task requirements, and sensory context (Niell and Stryker 2010; Harris and Thiele 2011; Reimer et al. 2014, 2016; McGinley, David, et al. 2015; McGinley, Vinck, et al. 2015). While QW has classically been studied as a state that qualitatively differs from active behavior, it has recently been shown to be a dynamic state in which transitions from low to high arousal are accompanied by increasing changes in cortical functioning (McGinley, David, et al. 2015; Neske et al. 2019). However, it is still unknown how these spontaneous changes in QW affect cortical dynamics during sensory processing. This study aims to elucidate whether spontaneous changes in the arousal state during QW are associated with specific brain signatures and whether these changes might affect visual stimulus processing.

Fluctuations in pupil diameter reflect changes in behavioral state and can occur spontaneously or evoked by stimuli. The Locus Coeruleus-Norepinephrine (LC-NE) system is regularly implicated as the neural substrate of stimulus-evoked pupil dilation, via changing arousal state, primarily because of the anatomical connections between the LC and the sympathetic and parasympathetic nervous system (Aston-Jones and Cohen 2005; Joshi et al. 2016; Joshi 2021). Furthermore, brief spontaneous dilations of the pupil diameter during QW correlate with desynchronization of the membrane potential of supragranular cortical neurons, and with an increase of low-frequency oscillations in neocortex (Reimer et al. 2014, 2016). Thus, continuous fluctuations in behavioral state can be separated according to spontaneous changes of the pupil diameter, and groups from both sides of this arousal distribution can be contrasted with each other to unveil differences in cortical dynamics during stimulus processing.

We used the ferret, an animal model with a visual cortex organized similar to primates (Kaschube et al. 2010), to investigate the neuronal correlates of arousal states during QW. We presented full-field and full-contrast gratings to awake, head-fixed ferrets and studied rhythmic changes in the local field potential (LFP) and spiking activity of neurons in ferret primary visual cortex (area 17, V1) and the posterior parietal caudal cortical area (PPc), a higher order sensory area that shows connections to visual cortical areas (Dell et al. 2019). We found that pre-stimulus pupil diameter spontaneously fluctuated during QW. We separated the distribution of pupil diameters into quintiles and compared the electrophysiological changes observed in the lowest and highest quintiles of the distribution. During short periods of pre-stimulus pupil dilation, corresponding to periods of high arousal during QW, we observed an increase in orientation tuning selectivity and spike-count correlations between spiking responses.

Furthermore, we observed amplitude and peak shifts of the spectral power of the LFP signal depending on the QW arousal state. The high-arousal quintile within QW states triggered low-frequency interareal increments in coherence, and computational models able to reproduce these effects pointed towards feedback signals from higher brain areas as a potential cause. The high-arousal condition in QW also led to frequency-dependent phase-locking of different neuronal types in the striate cortex. Our results show that the variability and sensitivity of cortical responses to a stimulus critically depend on the animal’s behavioral state before stimulus onset.

## Materials and Methods

All animal experiments were conducted with the approval of the local ethical committee of the University of Amsterdam and the Netherlands National Committee for the protection of animals used for scientific purposes. We used six healthy, adult female ferrets (*Mustela putorius furo*) of approximately six months old at the onset of the study. Ferrets were group-housed, maintained under a 16/8 hours dark/light cycle, and received water and food *ad libitum*.

### Stimulation paradigm

We used custom software coded in MATLAB (*Mathworks*) and the Psychophysics Toolbox (Brainard 1997) to present visual stimuli to awake ferrets. All stimuli were presented on an LCD monitor (53.3x33 cm, refresh rate 60 Hz). The ferrets were positioned on an elevated platform inside a sound-attenuated Faraday cage during the electrophysiological recordings, and habituated to sitting comfortably in a custom-made cylindrical body holder (Figure 1A). We placed the monitor at ∼25 cm in front of the animals (Figure 1A). Visual stimuli consisted of presenting a whole field chevron pattern (SF = 0.18 cycles/deg, TF = 1.2 deg/s) for 1 second in eight different orientations (0° to 315° in steps of 45°), interspersed with an intertrial interval (ITI) segment of an isoluminant gray screen, shown for 0.35 s with a random jitter of ±20 ms (Figure 1B). Following 11 stimulus repetitions, we presented another isoluminant gray screen for 3 s to record baseline activity. We tracked eye position and pupil diameter based on the corneal reflection of light with a non-invasive monocular eye tracker (ISCAN ETL-200, Rodent Eye Tracking Lab).

**Figure 1.**
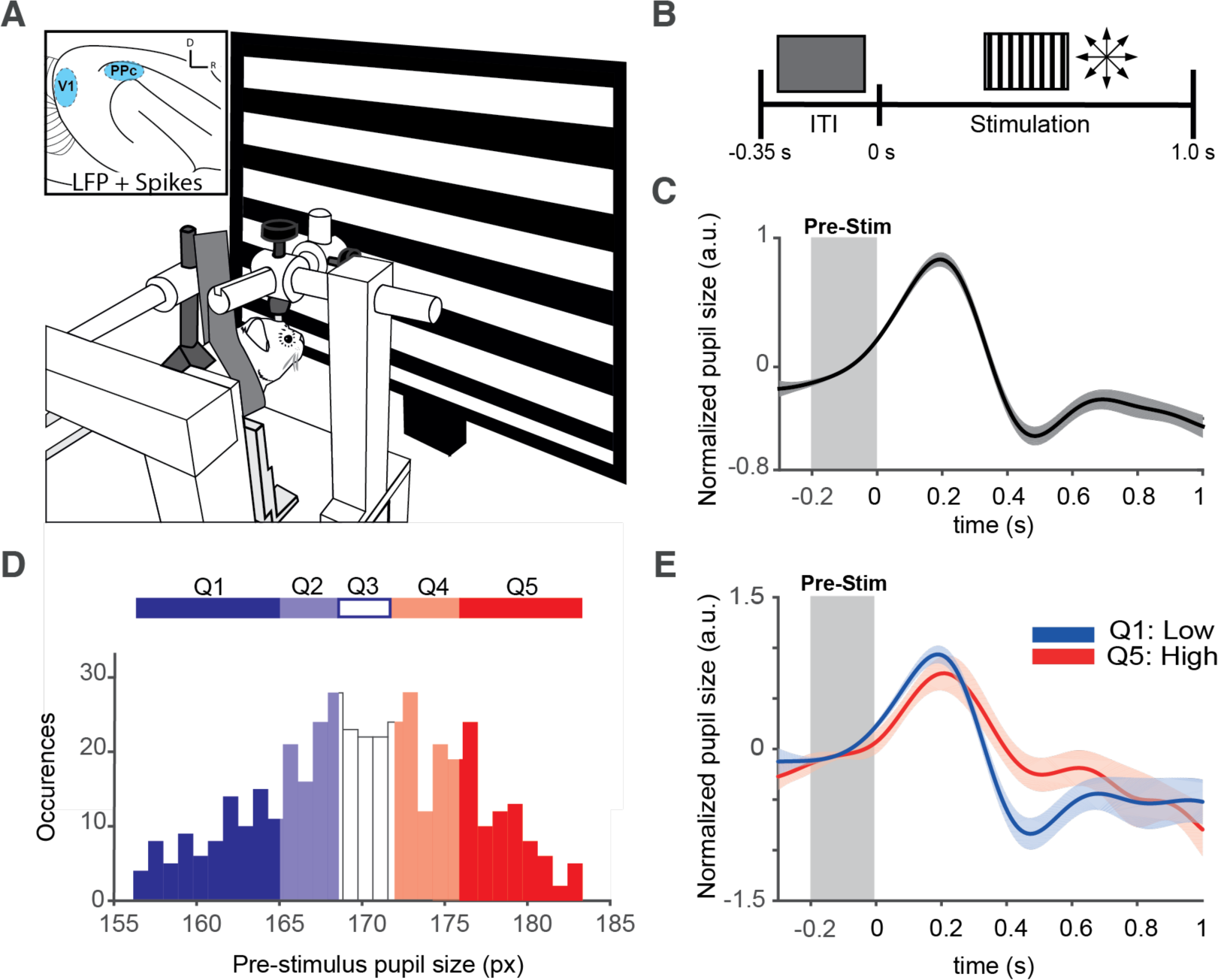
Spontaneous pupil fluctuations during passive visual stimulation. **A)** Experimental design. Awake head-fixed ferrets were placed in a cylindrical holder, to which they were familiarized, in front of a computer screen displaying visual stimuli. Inset: a schematic of the ferret cortex displaying the target areas for recordings (V1 and PPc). **B)** The experiment consisted of the presentation for 1 s of a drifting grating (SF=0.18 cycles/deg, TF=1.2 deg/s) for eight different orientations. During the inter-trial interval (ITI), a grey screen was presented for 0.35 s with a SOA of ± 0.02 s. **C)** Pupil response of one ferret normalized over all trials. For visualization purposes, pupil trials were smoothed with a low-pass Butterworth filter with a cut-off frequency of 3 Hz and subsequently averaged across all ferrets and sessions. The onset of the stimulus occurs at 0 s. The period of 0.2 s before stimulus onset (in gray) corresponds to the pre-stimulus control window. **D)** Pre-stimulus pupil sizes (expressed as number of pixels, *px*) were divided into quintiles. **E)** Pupil responses for the first quintile (Q1, low arousal) and the last quintile (Q5, high arousal) normalized per quintile. The normalized data shows a similar pupil dynamic range for both conditions (Joshi 2021).

### Headpost implantation

To maintain a stable visual field throughout recordings, ferrets received a cranial headpost-implant in two separate surgeries. First, ferrets were implanted with a titanium baseplate under aseptic and sterile surgical conditions. The baseplate was screwed to the skull’s frontal bone with four titanium screws and fortified using C&B Super Bond (Parkell) under general anesthesia. We used Ketamine (10 mg/kg) and Dexdomitor (0.08 mg/kg) administered i.m. to induce general anesthesia, and isoflurane (1–3%) delivered in 100% medical-grade O2 for maintenance. Marcaine 0.5% was used as a local anesthetic for intubation. We used Buprenorphine (0.01 mg/kg, s.c., pre-operative) and Meloxicam (0.2 mg/kg, s.c., peri- and post-operative) as analgesia, and Amoxicillin (20 mg/kg, s.c., in a single-dose peri- and post-operative) for antibiotic prophylaxis. Atropine (0.05 mg/kg, s.c.) and Dopram (5 mg/kg, s.c.) were used in case of respiratory distress. Dehydration was prevented using 50 ml of saline (0.9%, s.c.). We routinely monitored the respiratory rate, heart rate, and internal body temperature during surgery to maintain physiological values (LifeVet M, Eickemeyer). We used Antisedan (0.8 mg/kg, i.m.) to reverse the action of Dexdomitor at the end of the surgery. After an average of four weeks of ossification around the baseplate’s anchor screws, we installed a head-post pole under light isoflurane anesthesia to allow head-fixed electrophysiological recordings. A small skin incision was performed above the baseplate, and the pole was attached and secured with Loctite 242. Meloxicam (0.2 mg/kg, s.c.) was used as post- operative analgesia.

### Electrode implantation

Under the same surgical conditions, we performed a 1.5 mm diameter craniotomy over the primary visual cortex of the ferret (*area 17, V1*) using cranial, gyral, and sulcal landmarks (Innocenti et al. 2002; Manger et al. 2004). Once the dura was exposed and retracted, a 32-channel multielectrode silicon probe (*Models A4x8-5mm-200-400-177-CM32 and A2x16-10mm- 100-500-177-CM32,* NeuroNexus) was inserted approximately 1.5 mm along the dorsoventral axis in the visual cortex (n=6). Half of these animals (n=3) received an additional implant of the same probe in the posterior parietal cortex (PPc). The electrophysiological implant was covered with a protective cap and chronically fixed with dental cement (Simplex Rapid Acrylic Powder, Kemdent). The wound was closed by suturing the skin or gluing with Vetbond (3M). The electrode placement was verified posthoc using a conventional Nissl staining procedure in brain slices of 100 μm thickness.

### Electrophysiological recording techniques and signal preprocessing

We recorded spiking and local field potential (LFP) activity in V1 and PPc using an analog *Neuralynx ERP-54* system and collected data with the *CheetahRev 5.6.3* (Hardware SubSystem Cheetah 64) software. The signals were recorded with an input range of 1506 µV and a sampling frequency of 30303 Hz in 32 Continuously Sampled Channels (CSC). The electrophysiological signals were amplified with an amplitude gain of 830 dB and filtered with a band-pass filter between 0.1 to 9000 Hz. Transistor–transistor logic (TTL) pulses for time stamping of stimulus presentation were serially sent through an Arduino Board Mega 2560 to the acquisition system. All signals were recorded continuously for the entire duration of the recording session. The recorded signals were processed offline using the FieldTrip toolbox (https://www.fieldtriptoolbox.org/) for electrophysiological analyses (Oostenveld et al. 2011) and MATLAB custom software. We used the open-source software *Kilosort2* to extract single-unit spikes from the broadband signal (Pachitariu et al. 2016). Following automatic spike detection, we performed an additional manual curation of those spikes ambiguously separated. The multi-unit activity was obtained from the broadband signal using a high-pass filter at 500 Hz (Butterworth filter, second-order). For LFP analyses, the original broadband signal was detrended, linearly downsampled to 1024 Hz, and low-pass filtered with a Butterworth filter (second-order) with a cut-off frequency at 100 Hz. We removed powerline artifacts from the LFP using a band-stop filter centered at 50 Hz and harmonics. Then, we segmented the continuous signals in epochs of interest, for each channel, between 0.3 s pre-stimulus onset to 1 s post-stimulus onset.

### Trial Inclusion criteria and grouping

We visually inspected the stimulus-evoked-pupil response of the eye-tracking recordings to assess the quality of the traces. We eliminated trials in which an accurate pupil size could not be determined during the presentation of visual stimuli or the pre-stimulus baseline. To determine the arousal state of the animal before stimulation and define the high and low arousal conditions, we selected a time epoch between -0.2 s and stimulus onset as a pre-stimulus baseline window. From this window, we obtained the distribution of the average pupil size. We then divided the pre-stimulus pupil size distribution into quintiles (Figure 1D), selecting the trials of the first and fifth quintiles for further analyses. We denominated those trials of the first quintile as the low arousal group (condition QW-Low arousal) and the trials of the fifth quintile as the high arousal group (condition QW-High arousal).

### Orientation Selectivity Index

We calculated an orientation selectivity index (OSI) to measure the orientation-tuning properties of the recorded neurons in ferret primary visual cortex. First, we computed the mean discharge rate (in spikes/s) for each stimulus orientation (8 in total) using a spike density function and convolving the spike trains with a gaussian kernel. Then, after identifying the orientation that evoked the highest rate over trials (Ratepref) and inferring the orthogonal orientation (Ratenon-pref), we computed the neuronal *d’* (Berens et al. 2008; Meijer et al. 2017, 2020) as an orientation selectivity index, a measure that includes the pooled variance across neurons, according to the formula:

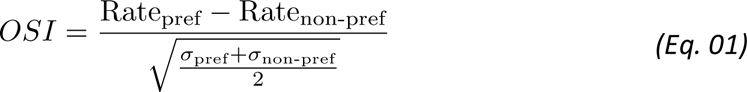

Following Meijer et al. (2017), we selected an OSI>0.4 as a cut-off value to classify neurons as orientation-selective.

### Spike-Count Correlations (rSC)

To obtain the pre- and post-stimulus correlated variability of spiking responses across high- and low-arousal conditions, we calculated the Pearson correlation of spike counts (noise correlations) for every pair of simultaneously recorded single units (Cohen and Kohn 2011) from ferret primary visual cortex. To remove the potential influence of confounding variables that could affect the variability of spiking responses, we transformed all spike counts from every neuron into a z-score using the mean and SD for every repetition of both experimental conditions (Nandy et al. 2017; Arbab et al. 2018). Then, we pooled these z-scored values in pairs of neurons and obtained the Pearson correlation from these pairs. The rSC was calculated for each condition using a spike counting window of 0.2 s across the entire epoch of interest (Figure 2B). We controlled the trial-to-trial variability in spike-count correlations by shuffling the neurons across conditions before calculating the rSC and repeating the analysis through different spike count windows, ranging from 0.05 to 0.3 s (Kass and Ventura 2006; Arbab et al. 2018).

**Figure 2.**
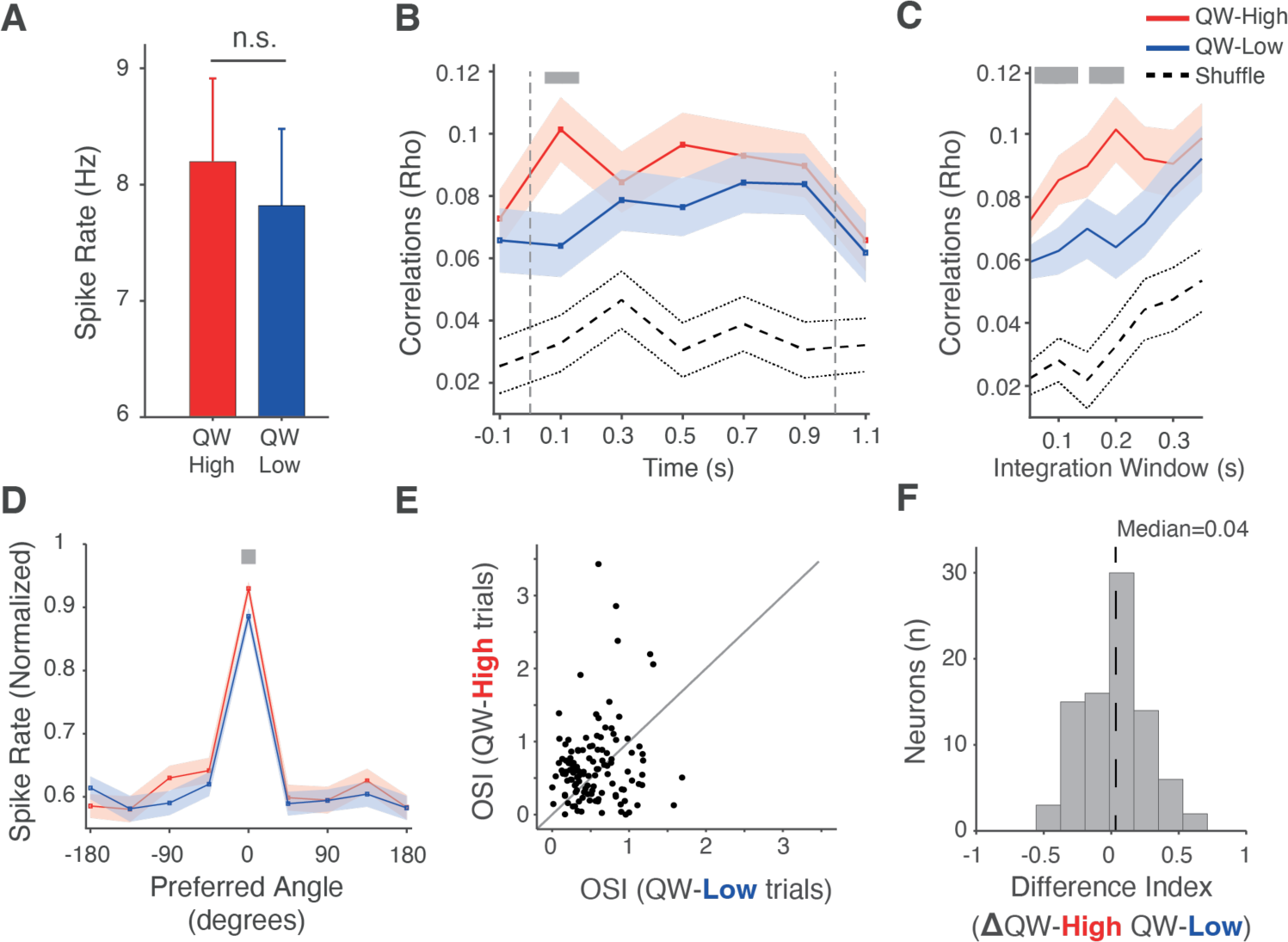
Neuronal responses in primary visual cortex during high- and low-arousal quiescent states. **A)** Average (± sem) spike rate in V1 across the stimulation period (0.8 s post-stimulus onset) across all channels, sessions, and ferrets. Differences are non-significant (QW-L: 7.9 ± 0.7 Hz, QW-H: 8.2 ± 0.7 Hz, p=0.86 rank-sum=2.1x10^4^, Wilcoxon rank-sum test). **B)** Spike-count correlations (rSC) as a function of time. The rSC values were calculated using an integration window of 0.2 s. The red and blue lines represent the average rSC values ± sem (ribbon) across recorded neurons for high- and low-QW arousal conditions, respectively. Gray vertical dashed lines at the beginning and end of the trial represent stimulus onset and offset, respectively. Black dashed lines represent the average (± sem) of rSC between neurons in which the condition labels were randomly shuffled. Grey bar denotes a p- value < 0.05 in a randomization test, corrected for multiple comparisons across observations. **C)** Spike- count correlations calculated at 0.1 s post-stimulus onset as a function of different temporal integration windows. Gray bar and black dashed line as in *(B)*. **D)** Normalized spike rate (average ± sem) as a function of the preferred orientation angle (in degrees). Grey bar denotes a p-value < 0.05 in a randomization test, corrected for multiple comparisons across observations. **E)** Scatter plot of the orientation selectivity index (OSI) values during high and low arousal during QW. Each dot represents the average values of a recorded neuron. OSI values significantly increased during pupil dilation (p=0.023, z=2.26, sign=48; Sign test). **F)** Histogram of the OSI differences between high and low QW. Positive values indicate a higher OSI during QW-high. The dashed line represents the median of the distribution (0.04). The distribution of differences was significantly distinct from zero (p = 0.03, z = 2.05, rank-sum = 8591; Wilcoxon rank-sum test).

### Spectral analysis of power

The raw LFP data from each channel was separated into two different groups (QW-high and QW-low arousal conditions) corresponding to the first and fifth quintile of the pupil size distribution across trials. The resulting epochs of interest were demeaned and divided by their standard deviation. From these segments, we used a 1 s segment of the baseline window (from 1.5 to 2.5 s) and a 1 s period from the stimulus onset, comprising the entire visual stimulation. Then, we obtained the spectrally decomposed Fourier coefficients of every epoch of interest (per channel), applying a discrete fast Fourier transform (FFT) to the segmented trials.

We used the irregular-resampling auto-spectral analysis (IRASA) method to obtain the spectral power, which isolates the oscillatory components from the fractal (1/f) activity of the power spectrum of any neurophysiological signal (Wen and Liu 2015). IRASA produces an estimation of the oscillatory and fractal power spectral components by resampling the neural signal using multiple non-integer positive numbers and their reciprocals. Then, it calculates the geometric mean of the auto-power spectra of each resampled signal. The resulting spectrum contains a redistribution of the fundamental and the harmonic oscillatory frequencies by an offset that varies with the resampling factor. Conversely, the fractal component of the spectral estimation remains constant independently of the resampling factor. Finally, we obtained an approximate estimate of the power spectrum of the oscillatory component by subtracting the median of the mean auto-power spectra (containing the fractal estimation of the power) to the original power spectrum (obtained from the previously calculated Hanning tapered Fourier coefficients). The power estimates were normalized per session and animal relative to the mean power between 4 and 100 Hz (Malkki et al. 2016).

### Phase-locking of LFP-LFP signals

The LFP-LFP phase-locking value across electrodes was calculated using the Weighted Phase Lag Index (WPLI), a bivariate metric of the phase consistency across signals, less affected by volume conduction, noise and sample size (Vinck et al. 2011). Specifically, the WPLI estimates, for a particular frequency, the non-equal probability of phase leads and lags of the imaginary part of the cross-spectrum, weighted by the magnitude of this imaginary part of the cross-spectrum. The WPLI was computed using the equation:

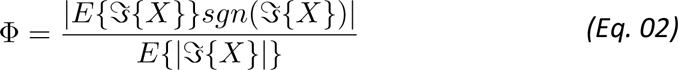

where *^sgn^*^(^*^={X}^*^)^ represents the signed estimation of the imaginary part of the cross-spectrum between two channels. The WPLI metric was estimated from a four-tapered multitaper Fourier decomposition (Jarvis and Mitra 2001) over an epoch of 1 s duration, resulting in a spectral smoothing of ± 2 Hz.

*Phase locking of spike-LFP signals.* The strength of the phase-locking between the LFP and the spikes was measured using the pairwise phase consistency analysis (PPC) (Vinck et al. 2012). For each frequency bin *f*, we determined spike-LFP phases in epochs of 5/*f* (5 cycles) length centered around the spikes obtained during the baseline and stimulation period. The PPC index was obtained from the Kaiser-tapered (*β*=3) Fourier coefficients and calculated according to the formula:

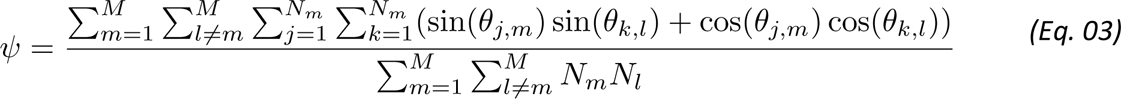

where *θj,m,* and *θk,l* are the *j*^th^ and *k*^th^ spikes at frequency *f* in trial *m* and trial *l,* respectively. *Nm* and *Nl* denote the number of spikes *N* in different trials. This method solves the problem of statistical dependence between spike-LFP phases since it restricts the PPC analysis to those spikes recorded from separate trials (Vinck et al. 2012). The PPC analysis was calculated using the *spiketriggeredspectrum* function (method: ppc1) from the FieldTrip toolbox (Oostenveld et al. 2011). The PPC index was calculated per each neuron individually. The resultant PPC spectra were averaged across all neurons.

### Computational Model

The computational model is based on our previous work (Mejias et al. 2016; Lindeman et al. 2021) and involves three levels of description: intra-laminar, inter-laminar, and inter-areal (Figure 5A). We consider two cortical regions at the inter-areal level: the primary visual cortex (area 17, V1) and the PPc. Each area consists of two cortical layers, or laminar modules, which represent the inter-laminar level and simulate the dynamics of superficial (2/3) and deep (5/6) layers, respectively. Each laminar module has one excitatory and one inhibitory population (e.g., intra- laminar level) modeled using firing rate dynamics. Their respective firing rates *r*_*E*_(*t*) and *r*_*I*_(*t*) are described by the following equations:

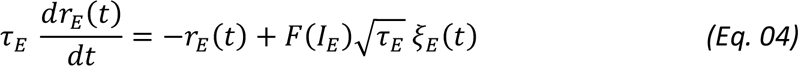

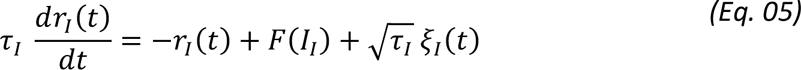

Here, τ_*E*_ and τ_*I*_are the time constants for the excitatory and inhibitory populations, respectively. ξ_*E*_(*t*) and ξ_*I*_(*t*) are Gaussian white noise terms of zero mean and standard deviation (σ) one. For superficial layers, we chose τ_*E*_ = 6 *ms*, τ_*I*_ = 15 *ms* for the time constants and σ = 0.3 for the noise strength, which leads to a noisy oscillatory dynamic in the gamma range. For deep layers, we chose τ_*E*_ = 42 ms, τ_*I*_ = 105 ms and σ = 0.45, which leads to a noisy oscillator in the alpha band frequency range. The relatively large values for the time constants in deep layers are thought to reflect other slow biophysical factors not explicitly included in the model, such as the dynamics of NMDA receptors.

The function *F*(*x*) = *x*/(1 − exp(−*x*)) is the input-output transfer function, applied for simplicity to all excitatory and inhibitory neuronal populations, which transforms the incoming input currents into their corresponding cell-averaged firing rates. The argument of the transfer function is the incoming current for each population, which involves (*i*) a background term, (*ii*) a local term, and (*iii*) a long-range term. The incoming current for excitatory and inhibitory populations, respectively, is:

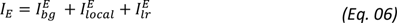

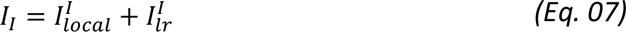

The background term is a default constant current only received by excitatory neurons in the visual cortex and area PPc and equals *I*^*E*^_*bg*_ = 3 for superficial excitatory neurons and *I*^*E*^_*bg*_ = 2 for deep excitatory neurons. The local term involves the input coming from neurons within the area, and is given by:

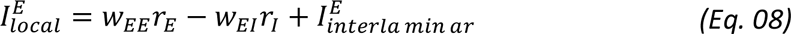

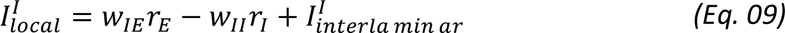

Here, wab are the synaptic weights from population b to a (*see Table 1 for numerical values*). The interlaminar terms represent contributions from a different layer than those terms used for the population under scrutiny. The only interlaminar projections are from superficial excitatory to deep excitatory neurons, with synaptic strength wds =1, and from deep excitatory to superficial inhibitory neurons, with strength wsd =0.75 (Mejias et al. 2016).

**Table 1:**
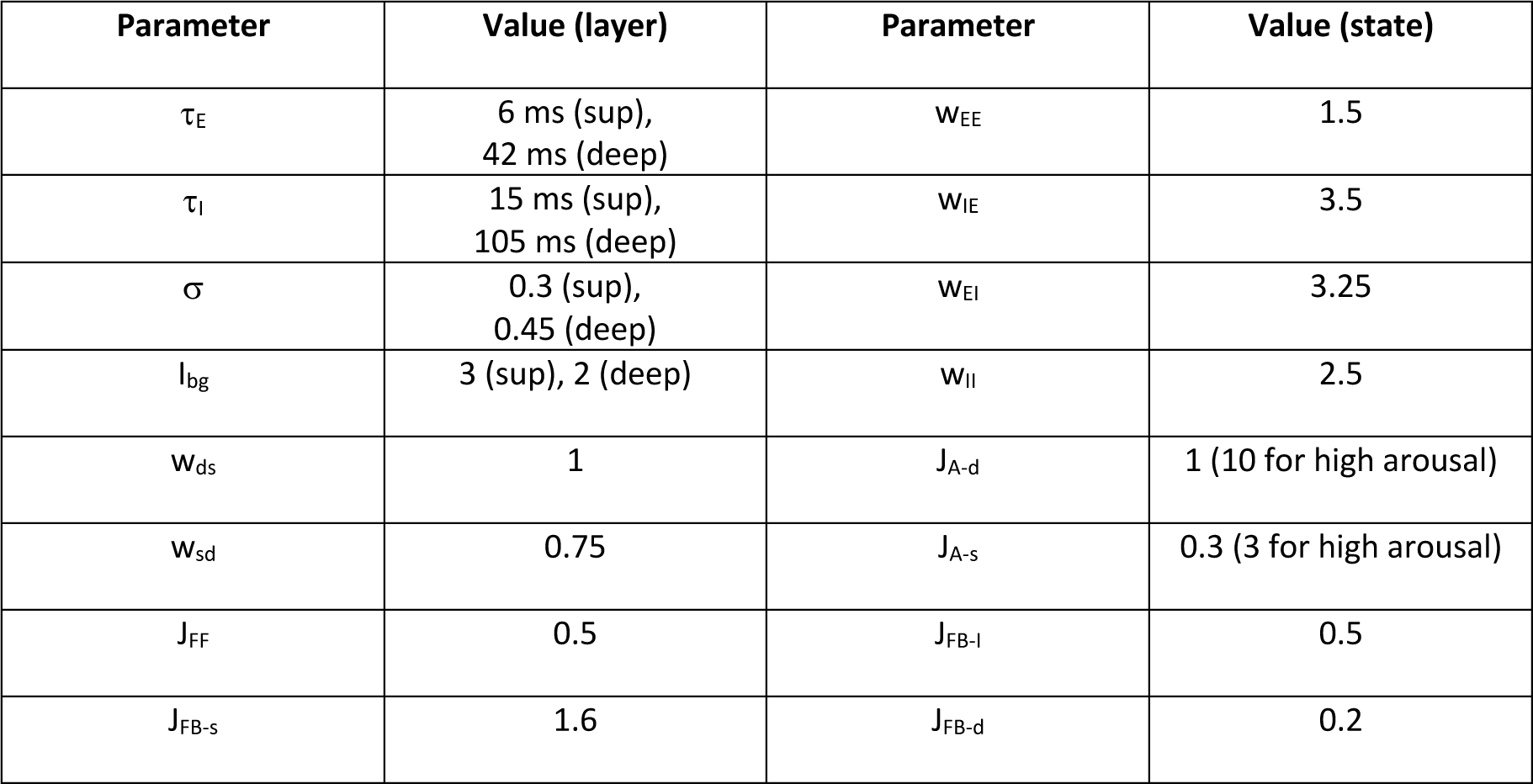
Parameter values of the computational model for V1-PPc laminar interactions.

| Parameter | Value (layer) | Parameter | Value (state) |
| --- | --- | --- | --- |
| $\tau_E$ | 6 ms (sup),<br>42 ms (deep) | $w_{EE}$ | 1.5 |
| $\tau_I$ | 15 ms (sup),<br>105 ms (deep) | $w_{IE}$ | 3.5 |
| $\sigma$ | 0.3 (sup),<br>0.45 (deep) | $w_{EI}$ | 3.25 |
| $I_{bg}$ | 3 (sup), 2 (deep) | $w_{II}$ | 2.5 |
| $w_{ds}$ | 1 | $J_{A-d}$ | 1 (10 for high arousal) |
| $w_{sd}$ | 0.75 | $J_{A-s}$ | 0.3 (3 for high arousal) |
| $J_{FF}$ | 0.5 | $J_{FB-I}$ | 0.5 |
| $J_{FB-s}$ | 1.6 | $J_{FB-d}$ | 0.2 |

With these parameters compatible with experimental values, the model produces noise-driven oscillators through stochastically perturbed stable focus dynamics − i.e. the activity behaves like a physical damped oscillator, which is kicked out of the fixed point by existing fluctuations. Furthermore, contrary to other models (Wilson and Cowan 1972), which utilize a limit cycle dynamic, the oscillatory activity produced here is weakly coherent from one cycle to another, providing a closer match to the highly irregular rhythmic activity of LFPs in actual neuronal recordings and reproducing a wide range of experimental findings (Mejias et al. 2016; Lindeman et al. 2021). Finally, the long-range term (*I*^*E*^_*lr*_ and *I*^**I**^_*lr*_ Eq. 6 and 7) in the input current includes currents coming from *J*_*ab*_*r*_*b*_ other neocortical area. These currents follow the general form *J*_*ab*_*r*_*b*_, (with *J*_*ab*_ being the synaptic strength from area *b* to area *a*), and therefore, we will specify only the synaptic coupling strengths to characterize them.

Following anatomical evidence widely consistent for mammals (Markov et al. 2014; Harris et al. 2019), we consider feedforward projections from primary visual to posterior parietal cortices originating in layer 2/3 pyramidal V1 neurons and target (indirectly via layer 4, which is not explicitly included in the model) pyramidal neurons in PPc layer 2/3 (with a synaptic strength of JFF = 0.5). Feedback projections stem from pyramidal neurons in deep layers of PPc and target pyramidal neurons in superficial (strength of JFB-s = 1.6) and deep layers (strength of JFB-d = 0.2), and inhibitory neurons in superficial and deep layers (strength of JFB-I = 0.5 for both cases). Here, arousal signals to these cortical areas are modeled as top-down feedback inputs arriving mostly in deep PPc layers (with a strength of JA-d = 1 for the low arousal case) and in superficial layers of V1 (strength of JA-s = 0.3 for low arousal, *see Table 1*), in agreement with recent hypotheses about the dependence of feedback targets on the hierarchical distance of the projections (Markov et al. 2014). For the high arousal signal, these top-down arousal inputs are multiplied by 10.

To mimic the depth of the recording electrodes for V1 and PPc in experiments, we estimate the LFP signal in the model by a weighted average of the excitatory superficial and deep layers, with a superficial-to-deep ratio of 2:8 for both areas (*i.e.,* 20% and 80% weight of the superficial and deep layers, respectively). These simulated signals were then used to compute spectral coherence (as in: Mejias et al. 2016) and the WPLI index.

### Statistical testing

We assessed significant differences between conditions using a nonparametric statistical approach (Wilcoxon rank-sum and sign tests) for the spike analyses with a significance threshold of p<0.05. For the spectral analyses, we calculated a nonparametric permutation test of the coherence differences (Maris et al. 2007), adjusted by a multiple comparison correction at the subject level (Scheeringa et al. 2011). First, the individual differences between electrodes (within-subjects) were computed using a Welch t-statistic. Next, we obtained individual p-values of the t-Welch differences from a Student’s t cumulative distribution function, using the Welch- Statterth-Waite equation to estimate the degrees of freedom of every distribution, and a cluster-based approach across frequencies to select significant frequency-band clusters (Maris and Oostenveld 2007). Then, we built a null distribution of these frequency bin clusters by computing a paired sample t-test and randomly exchanging the condition labels between conditions. We repeated this step across 1000 permutations of the data retaining the maximum and minimum values of the cluster-selected values. Finally, we compared the observed T-statistics to the maximal and minimum distributions. T-statistics were considered significant at p<0.05 if they were below the 2.5^th^ percentile of the minimum or above the 97.5^th^ percentile of the maximal distribution. We used a generalized linear mixed model (GLMM) to evaluate the power changes due to arousal across areas considering the variability across animals and channels as random effects (Tuerlinckx et al. 2006; Johnson et al. 2015). The statistical significance of the GLMM was tested using a likelihood ratio test under a χ^2^ distribution with a significant threshold set at p<0.05.

## Results

To investigate the effects of spontaneous arousal changes during QW on cortical microcircuits, we recorded the LFP signal and spiking activity from ferret primary visual cortex and the caudal part of the posterior parietal cortex (PPc) from six ferrets. Ferrets passively viewed full-screen gratings at eight different orientations for one second, while we continuously recorded pupil diameter changes used to infer behavioral state (Figures 1A and 1B).

### Pupil diameter spontaneously fluctuates during quiet wakefulness

We first focused on the pupil response of the animal during stimulus presentation (Figure 1C for the normalized pupil trace and Supplementary Figure 1A for the non-normalized pupil trace). As expected, pupil illumination produces a typical event-related response consisting of pupil dilation followed by a constriction that undershoots the pre-stimulus size period (Wang and Munoz 2015; Wang et al. 2017). Next, we wondered whether this event-related pupil fluctuation could be affected by the behavioral state of the ferret. We calculated the area of the pupil (in pixels) in a pre-stimulus window between 0.2 s and 0 s pre-stimulus onset (Figure 1C, Supplementary Figure 1A), using the average of such window as an indicator of the animal’s behavioral state before the stimulus appeared (Aston-Jones and Cohen 2005; Joshi et al. 2016; Joshi and Gold 2020; Joshi 2021).

Then, we used the average of this pre-stimulus window to create a pupil size distribution that we separated into quintiles (Figure 1D). Based on the distribution of pupil sizes, we considered those trials from the first quintile (Q1) exhibiting maximum constriction as low arousal trials. Conversely, the last quintile (Q5) trials displaying maximum dilation were considered high arousal trials. Since both quintiles pertain to a distribution of a QW state, we denominate both conditions as QW-low (QW-L) and QW-high (QW-H) arousal, respectively. In total, we used ∼80 trials per session per condition for subsequent analyses.

The normalized event-related pupil response of both conditions (Figure 1E, and Supplementary Figure 1B) shows a similar dynamic for QW-high and low arousal trials. In both conditions, the event-related pupil response consists of a post-stimulus pupil dilation followed by a pupil constriction. This pupil constriction is more prominent during low arousal, but the similar shape of the response suggests that the pupil diameter spontaneously fluctuates during passive stimuli similarly for low and high arousal conditions.

### High arousal increases the selectivity of spiking responses during quiet wakefulness

Active behavioral states improve neuronal coding efficiency in area V1 of primates (Shadlen and Newsome 1998; Mitchell et al. 2007; Cohen and Maunsell 2009; Ecker et al. 2010; Nandy et al. 2017; Denfield et al. 2018) and mice (Niell and Stryker 2010; Keller et al. 2012; Reimer et al. 2014, 2016; Vinck et al. 2015; Leinweber et al. 2017), but it is unknown whether encoding improvement can be observed during quiescence. Therefore, we investigated whether high and low arousal QW states contribute differently to the coding efficiency of cortical neurons. To answer this question, we recorded 145 single units from the primary visual cortex while ferrets passively observed eight different orientations of full-contrast gratings for one second per trial. From these 145, 110 neurons (75.8% of the total) responded to the presentation of visual stimuli. We checked whether different QW states change the spike rate of neurons responding to the stimulus presentation by computing an average spike rate across the stimulus presentation window. Previous studies have shown a decrease in the spiking rate of high-arousal quiescent periods following active locomotion, but the spiking rate within quiescent periods remains unaffected (McGinley, Vinck, et al. 2015). In agreement with these results, we observed that the spike rate of striate cortex neurons in different QW states does not change with passive stimulation (Figure 2A, QW-L: 7.9±0.7 Hz, QW-H: 8.2±0.7 Hz, p = 0.86, Wilcoxon rank-sum test: rank-sum = 2.1x10^4^).

We next focused on the correlated variability across neurons. An increase of correlated variability in a neuronal population is usually associated with decreased quality of the information represented in a population response (Shadlen and Newsome 1998; Averbeck and Lee 2006; Mitchell et al. 2009; Cohen and Kohn 2011; Arbab et al. 2018) (but see: Montijn et al. 2016) and can be modulated by active locomotion and attentional state (Reynolds et al. 2000; Cohen and Maunsell 2009; Mitchell et al. 2009; Reimer et al. 2014; Vinck et al. 2015). Therefore, we evaluated the response variability between the pairs of neurons during QW using spike-count correlations (rSC) between primary visual cortex neurons with an integration window of 0.2 s (Figure 2B). These correlations reflect the functional state of neuronal networks across time (Cohen and Kohn 2011). After stimulus onset, both conditions showed rSC values above chance level, indicating significant correlations. In addition, high arousal QW states showed a transient increase of rSC approximately at 0.1 s after stimulus onset (p<0.05, nonparametric randomization test across neurons). Under both conditions, the rSC slowly rose across time after 0.1 s post-stimulus onset, decreasing to baseline levels after stimulus offset. Finally, we controlled for bin size effects calculating the rSC centered at 0.1 s post-stimulus onset with integration windows of different sizes (Figure 2C). In all these cases, QW high-arousal states showed higher significant spike-count correlations than QW low arousal states (p<0.05, nonparametric randomization test across neurons), indicating that state fluctuations within QW can lead to different intra-areal spike correlations. Notably, our results show that the stimulus onset during quiescent states can trigger an increase of rSC despite high-arousal, suggesting, contrary to earlier findings (Shadlen and Newsome 1998; Averbeck and Lee 2003; Mitchell et al. 2009; Arbab et al. 2018), that a rise in spike-count correlations might contribute to improving the information processing efficiency during periods of increased arousal.

Previous studies have shown that active states effectively increase the orientation selectivity of striate neurons (Niell and Stryker 2010; Reimer et al. 2014; Vinck et al. 2015). For this reason, we evaluated whether high and low arousal QW states modulate the selectivity of the response of visual cortex neurons to drifting gratings using an orientation selectivity index (OSI), which considers the pooled response variability from different neurons across preferred and non-preferred orientations (*see Methods*). We observed that the spike rate on neurons at their preferred orientation was modestly but significantly higher during high arousal (Figure 2D, p < 0.05, nonparametric randomization test across neurons). Furthermore, individual OSI values significantly increased for all recorded neurons during pupil dilation (Figure 2E, p = 0.023, Sign test: sign differences = 48, z = 2.26). From 142 recorded neurons, 53% showed an increment of the OSI during QW-H. This increase was observed using an index of the difference across conditions of very selective neurons with high OSI values (Figure 2F). In this analysis, positive values of the index indicate an increase of the OSI during high arousal. Furthermore, the difference index distribution was significantly distinct from zero (p=0.03, Wilcoxon rank-sum test: z = 2.05, rank-sum = 8591).

In sum, these results show that high-arousal QW behavioral states increase the orientation selectivity index and the spike count correlation of the neuronal population. An increase of rSC values implies an increase in the synchronized activity of neurons due to common input. Furthermore, high rSC has been interpreted as a decrease in the quality of the information processing (Cohen and Kohn 2011). Here we showed that, despite high rSC values, QW states improve the response selectivity of neurons in V1 during stimulus processing.

### Quiet wakefulness induces amplitude and peak shifts of the spectral power

The local field potential (LFP) signal is a prime candidate to study the relationship between behavioral states and changes in cortical microcircuits functions. Rhythmic fluctuations of the LFP across time primarily reflect synchronized excitatory and inhibitory interactions among cortical microcircuits neurons (Buzsáki et al. 2012; Pesaran et al. 2018). Furthermore, these LFP oscillations have been associated with several cognitive functions and computational mechanisms (Fries 2009; Bosman et al. 2014; Singer 2018). Since behavioral states influence the synaptic activity of cortical microcircuits, we explored LFP power dynamics across different QW conditions during passive stimulation (Figure 3). We recorded the LFP activity of one primary sensory area (V1) and a hierarchically superior area (area PPc), focusing on the power changes between high vs. low-arousal quiescent periods in which animals passively observed the displayed gratings. We compared these power fluctuations with those elicited during a baseline period with no stimulation.

**Figure 3.**
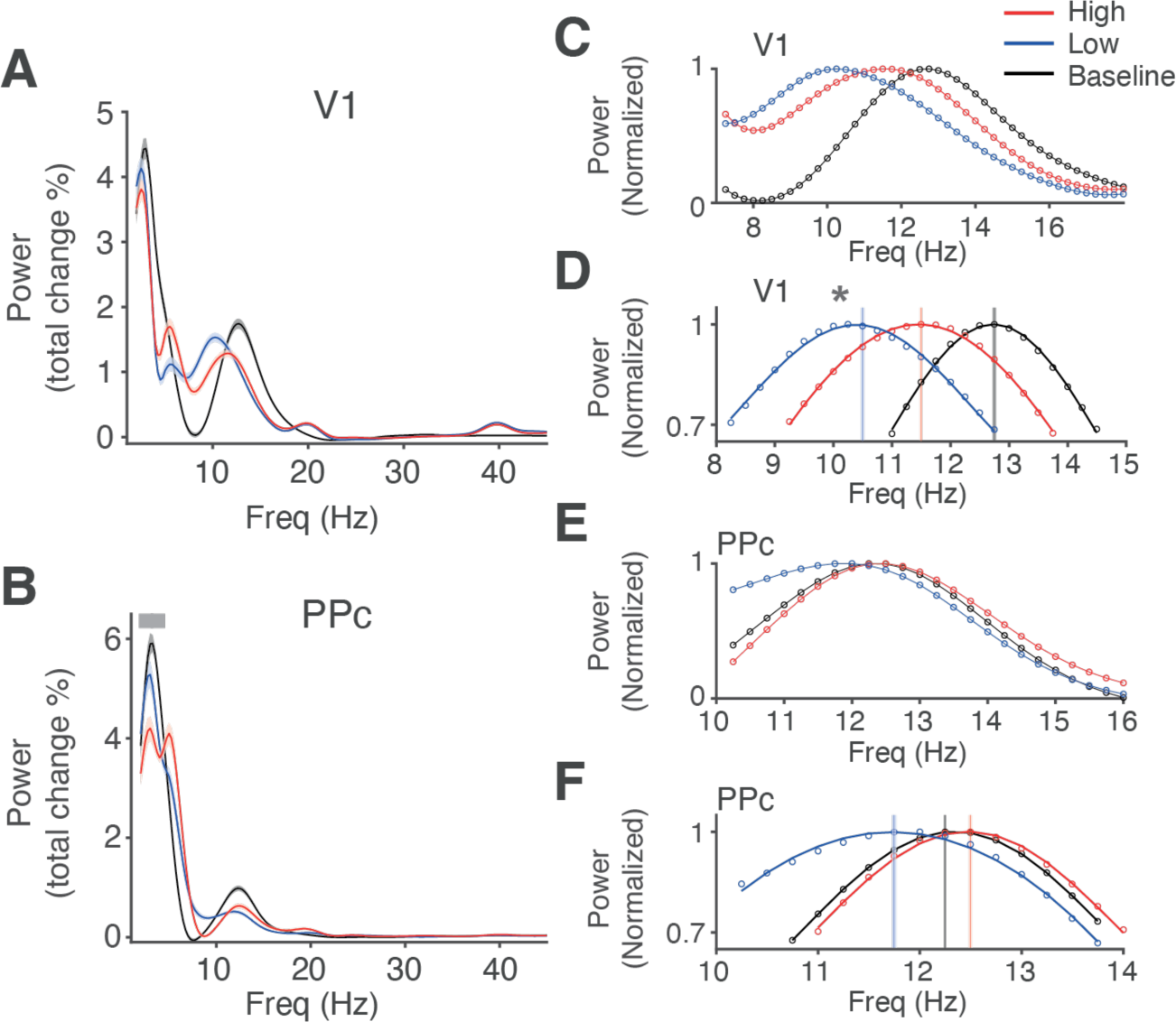
LFP Power estimation in primary visual and posterior parietal cortex during high- and low-arousal trials. **A)** Average power spectrum (% of the total power) across all channels, sessions, and ferrets in V1. Black, red and blue traces (average ± sem) correspond to the activity during baseline, high- and low-arousal conditions. The frequency cut-off of the IRASA power estimation was set at 50 Hz. **B)** Same as *(A)* but for posterior parietal cortex. The gray bar shows a significant decrease of the power at a frequency band between 1-4 Hz modulated by arousal (p=0.036, χ^2^(2)=6.65, likelihood ratio test). **C)** Individual power peaks of an 8 to 16 Hz frequency band in V1 for baseline (black), high- (red), and low- (blue) arousal conditions. Power values are normalized to the maximum. Scaled spectra are shown in dots. **D)** A cosine function was fitted to the top third of the power for each condition (R>0.98 for each curve). Vertical lines represent the peak of the frequency band. Shaded regions correspond to the 95% CI. The gray asterisk denotes a significant shift change due to arousal. (p=0.009, χ^2^(2)=9.25, likelihood ratio test. Frequency peaks: Baseline: 12.76 Hz, High arousal: 11.24 Hz, shift from baseline -1.51±0.47 Hz, Low arousal: 10.29 Hz, shift from baseline -2.47±0.55 Hz). **E)** Same as *(C)* but for posterior parietal cortex. **F)** Same as *(D)* but for posterior parietal cortex. The GLM analysis did not show statistical significance. (p=0.07, χ^2^(2)=5.61, likelihood ratio test. Frequency peaks: Baseline: 12.25 Hz, High arousal: 12.51 Hz, shift from Baseline: 0.26±1.13 Hz, Low arousal: 11.78 Hz, shift from baseline -0.47±1.19 Hz).

We observed several changes associated with the animal’s behavioral state (Figures 3A and 3B). In the visual cortex, visual stimuli induced two narrow frequency band activities centered at 20 and 40 Hz, and a decrease in power amplitude between 1-4 Hz (Figure 3A). In area PPc, we also observed a decrease of the power between 1 and 4 Hz after the presentation of a stimulus. Importantly, the power amplitude at the peak of this frequency band shows a modest but significant decrease by about 2±0.7% of the total power in area PPc during high arousal QW (Figure 3B, p=0.036, χ^2^(2)=6.65, likelihood ratio test). Our results indicate that stimulus induced power changes at PPc are modulated by the state of arousal during quiescent wakefulness.

In addition, we observed a peak shift within an 8 to 16 Hz frequency band as a function of the arousal state in V1, but not in area PPc (Figure 3C to 3F). To estimate the peak value for each condition and area, we scaled the spectra and fitted a cosine to the upper third of the band (Figure 3D and 3F, all R values > 0.99), and we used a mixed model with the arousal conditions as a predictor to statistically evaluate this spectral shift. In V1, visual stimulation induced a significant shift of the power peak towards lower frequencies. In addition, low arousal shifted the peak towards lower frequencies compared with high arousal (see Figure 3C for the entire power spectrum comparison and 3D for the peak differences; χ^2^(2)=9.25 p=0.009, likelihood ratio test. Frequency peaks per group: Baseline: 12.76 Hz; High arousal: 11.24 Hz, shift from baseline -1.51±0.47 Hz ; Low arousal: 10.29 Hz, shift from baseline -2.47±0.55 Hz). In contrast, neither visual stimulation nor QW states induced significant power peak changes in area PPc (Figure 3E and 3F; χ^2^(2)=5.61 p=0.07, likelihood ratio test. Frequency peaks: Baseline: 12.25 Hz, High arousal: 12.51 Hz, shift from Baseline: 0.26±1.13 Hz, Low arousal: 11.78 Hz, shift from baseline -0.47±1.19 Hz).

### High- and Low arousal conditions are differentially coupled to frequency-dependent interareal coherence

Behavioral states are considered global phenomena (Harris and Thiele 2011) with uniform neural underpinnings across different brain areas. However, previous studies have stressed that functional connectivity at macroscopic and microscopic levels depends on both anatomical connectivity and behavioral state (Crochet et al. 2006; Poulet and Petersen 2008; Olcese et al. 2016, 2018; Poulet and Crochet 2019). Our power amplitude analyses suggest that low and high arousal states during QW elicit different power frequency band signatures in visual and parietal cortices. Therefore, we wondered whether visual and parietal cortices show similar or different functional connectivity profiles during QW. We used the WPLI index, which measures the phase consistency across two oscillatory signals, correcting for volume conduction (*see Methods*), as an estimate of the functional communication within and between visual and parietal channels (Figure 4, all significant results obtained with p<0.05 using a nonparametric randomization test corrected by multiple comparisons. *See Methods*).

**Figure 4.**
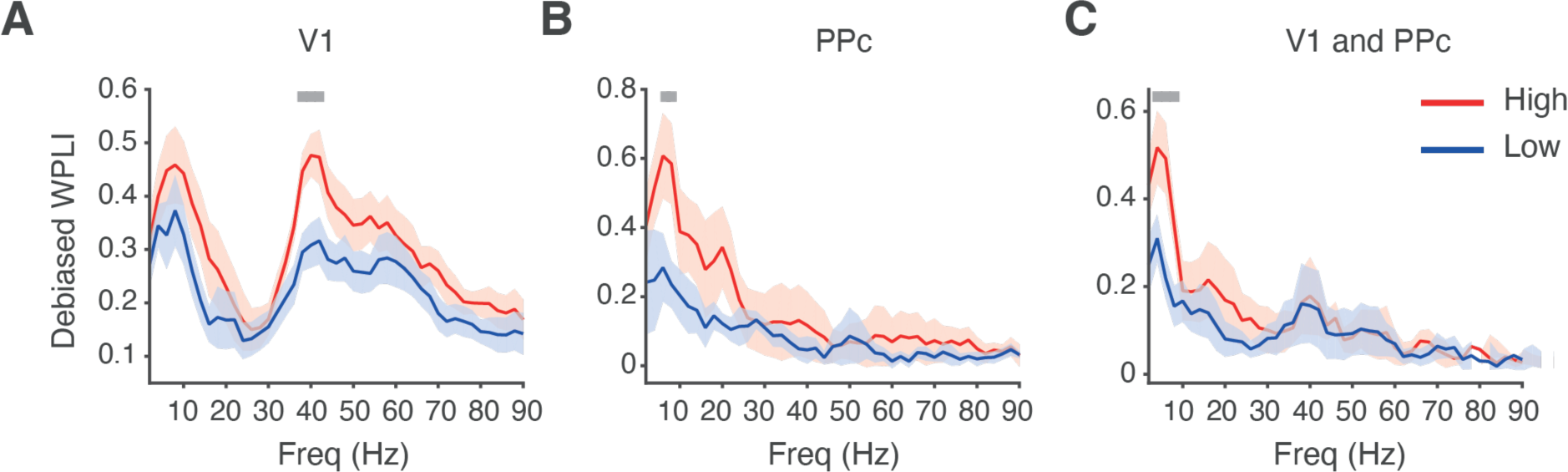
Intra- and interareal coherence between primary visual and parietal cortices during visual stimulation increases during a high-arousal quiescent state. A) Weighted phase lag index (squared WPLI debiased, a measure for LFP-LFP phase-locking connectivity, *see Methods*) for all channel combinations in V1 (average ± sem) and high- (red) and low (blue) arousal conditions. The gray bar shows a p < 0.05 level corrected for multiple comparisons across frequencies (nonparametric randomization test across site pairs; *see Methods*). A significant increase of the WPLI index, centered at a narrow gamma frequency band (40-50 Hz), is observed during high-arousal quiescent states. B) Same as *(A)* but for channel pairs within the posterior parietal cortex. C) Same as *(A)* but for channel pairs between V1 and posterior parietal cortex. In (B) and (C), a significant increase of the WPLI, centered around a theta frequency band (4 to 8 Hz), is observed during high-arousal quiescent states.

The average WPLI spectrum across channels within the visual cortex showed a significantly increased narrow band centered at 40 Hz during high arousal relative to low arousal (Figure 4A). Conversely, we observed a significant increase of the WPLI at 4-8 Hz in the parietal cortex during high arousal (Figure 4B). When we evaluated the WPLI spectrum across areas, we observed that a similar 4-8 Hz increase during high arousal dominated functional connectivity between areas (Figure 4C).

Our WPLI results suggest that functional connectivity increases during high arousal but with a different spectral profile depending on the area. Enhanced low-frequency WPLI is observed within PPC and between PPC and the visual cortex. In contrast, WPLI within V1 showed an increase in gamma-band synchronization. Our results show that the functional connectivity between brain areas in the ferret resembles that observed in the visual cortical system of non-human primates (Kerkoerle et al. 2014; Bastos et al. 2015), possibly reflecting hierarchical relationships between brain regions (Markov et al. 2014).

### Simulation of functional connectivity patterns by a computational model

Inter and intra-area anatomical connections carry feedback and feedforward information targeting specific laminar compartments, depending on the hierarchical level of sending and receiving areas (Markov et al. 2013). We wondered whether the observed differences in connectivity between high and low arousal states during QW can be simulated by differentially modulating the activity of the cortical laminae. Therefore, we modified a previously developed large-scale computational model constrained by weighted connectivity data derived from the macaque cortex (Mejias et al. 2016), testing different feedback connectivity profiles between V1 and PPc (Figures 5A and 6, see *Methods* for a model description).

**Figure 5.**
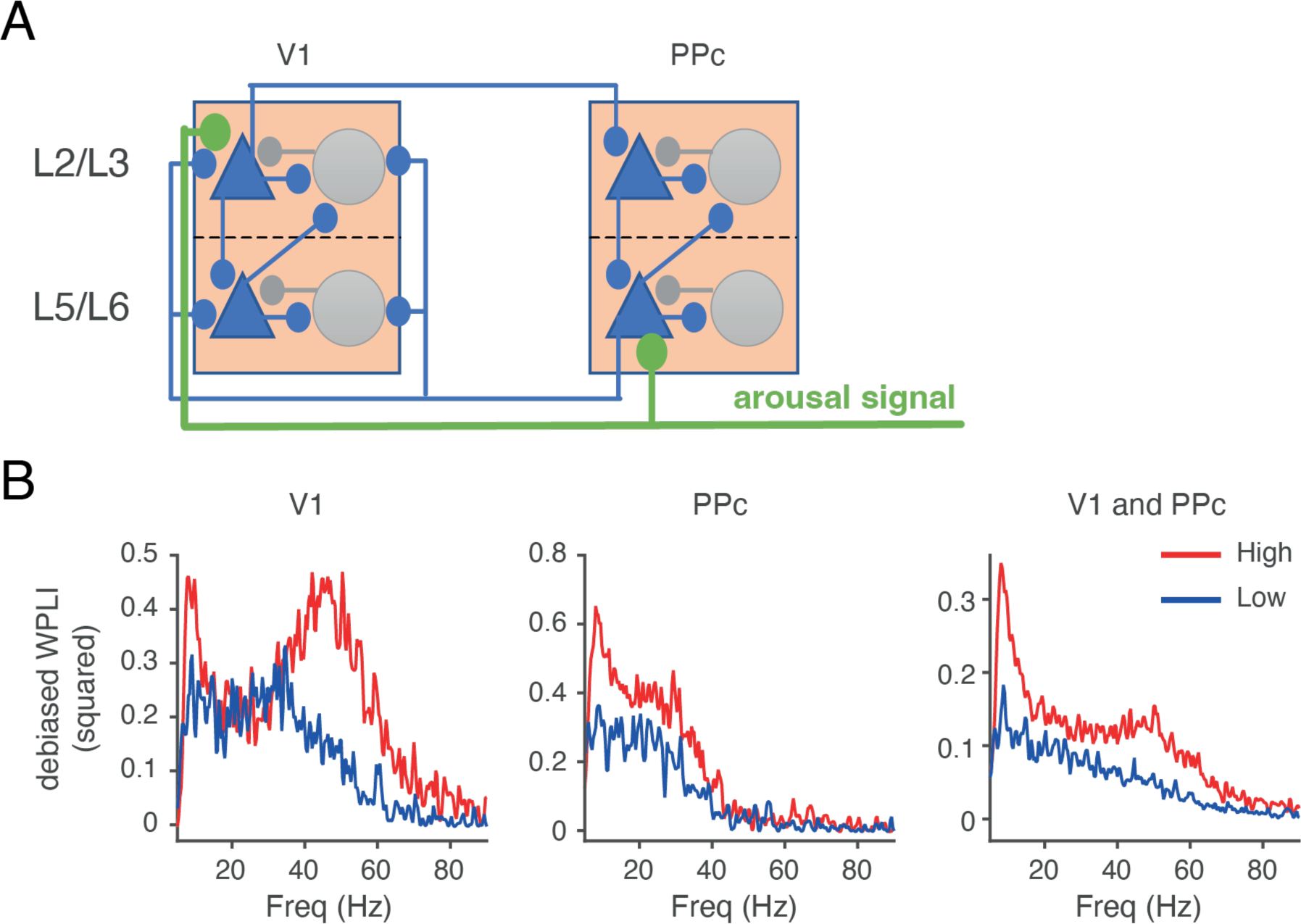
A computational resembles the WPLI coherence observed between V1 and PPc. **A)** Schema of the interlaminar circuit used to model the results observed in *Figure 4*. A simplified, minimal model or a cortical column is used to describe the laminar-specific interactions between V1 and PPc and the influence of feedback (FB) signals on them. In green, top-down projections carry arousal signals to the supragranular compartment of V1 and the infragranular compartment of PPc. **B)** Computational prediction of the functional connectivity, measured as WPLI-debiased index, within area V1 (left panel), within area PPc (middle panel), and between area V1 and PPc (right panel). Color codes are the same as in *Figure 4*. The configuration depicted in *Figure 5A* approximates the results observed in *Figure 4 (see* Table 1 *for the model parameters)*.

**Figure 6.**
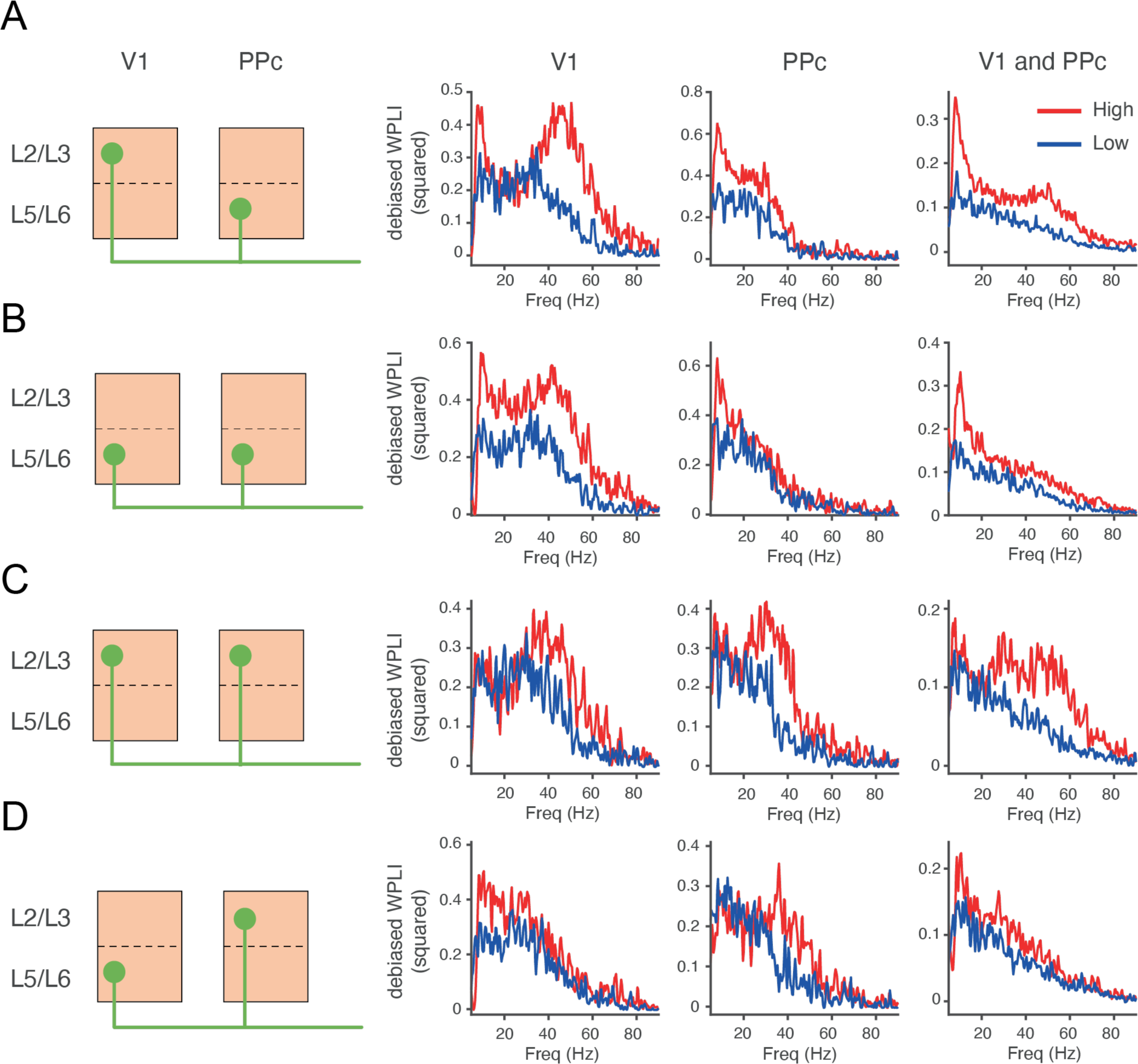
Inter and intra-area coherence estimates vary according to the pattern of interlaminar projections. **A to D)** *Left:* Variations in interlaminar feedback projection patterns used to test the functional connectivity model. *Right:* WPLI spectra of the functional connectivity within and between primary visual cortex and PPc (plotting conventions as in *Figure 4*). Panel A is the same as used in *Figure 5*.

Figures 5B shows the WPLI index estimation obtained from the model architecture depicted in Figure 5A. Supplementary Figure 2 shows the coherence spectra estimates of the model. These coherence results constitute a model prediction for future studies, while results from WPLI can be directly compared with our existing data. Our assumption is that arousal signals are conveyed by excitatory feedback projections targeting the infragranular layers of PPc and supragranular layers of V1, thus allowing the replication of the abovementioned functional connectivity results. Our model establishes layer-specific feedback modulations as a potential mechanistic implementation of arousal signals to the visual and parietal cortex during different quiescent states. Importantly, the model does not inform us about the origin of such modulatory signals, which can be traced back to subcortical neuromodulatory signals or cortico-cortical feedback projections, for example (*see Discussion*).

We have also studied the predictions of our model when other types of laminar feedback projections are considered. Figure 6 compares four different connectivity patterns between infra- and supragranular layers and the estimated WPLI spectra for intra- and inter-area connectivity. Figure 6A duplicates the model from Figure 5 and compares it with three alternative scenarios with different ratios for infra/supra granular feedback intensities (Figure 6B to 6D, see Supplementary Figure 3 for the coherence spectra estimates of the same models). These alternative models resemble some but not all features of the observed data. For example, the alternative model offered in Figure 6B (i.e., feedback preferentially targeting deep layers of both cortical areas) provides a plausible alternative that shows a reduced gamma peak in V1-PPC interactions under low-arousal conditions (in line with experimental data), although at the expense of an increased alpha peak for V1 (not found in the data). Alternative models are shown in Figure 6C (feedback to superficial layers), and D (feedback to superficial layers of PPC and deep layers of V1). Importantly, these models cannot reproduce the observed results and are therefore less plausible explanations. A balanced feedback modulation through excitatory projections to supragranular and infragranular layers in primary visual and PPc cortices, respectively, is thus best consistent with our experimental findings on functional connectivity during high- and low-arousal quiescent.

In sum, our model establishes layer-specific feedback as a possible mechanistic implementation of arousal latter signals to the visual and parietal cortex. Furthermore, the feedback structure considered in this model is compatible with current hypotheses about the hierarchical distance of feedback generation signals and targeting downstream areas (Markov et al. 2013, 2014; Harris et al. 2019), suggesting a potential parallelism between feedback modulations in ferrets and macaques.

### Narrow and Broad Spiking V1 cells phase-lock to different frequency bands during quiescent high- arousal conditions

Since excitatory and inhibitory neurons entrain to local rhythms at distinct phases of the oscillatory period, with functional implications for cortical microcircuits (Hasenstaub et al. 2005; Siegle et al. 2014; McGinley, Vinck, et al. 2015; Vinck et al. 2016), we evaluated whether different neuronal populations specifically phase-lock to the observed LFP rhythms during high and low QW conditions. First, we sorted the 110 visually responsive neurons in V1 using the peak-to-trough duration of their waveform (Figure 7A) (Mitchell et al. 2007; Lansink et al. 2010; Vinck et al. 2016; Arbab et al. 2018). The distribution of peak-to-trough values was significantly bimodal (p=0.03, Hartigan’s dip test: dip=0.04). Then, we selected narrow-spiking and broad-spiking cells using 350 µs of the peak-to-trough duration as a criterion for separation. Using this procedure, we obtained 17 narrow-spiking and 84 broad-spiking cells. Nine neurons remained unclassified (Figure 7B).

**Figure 7.**
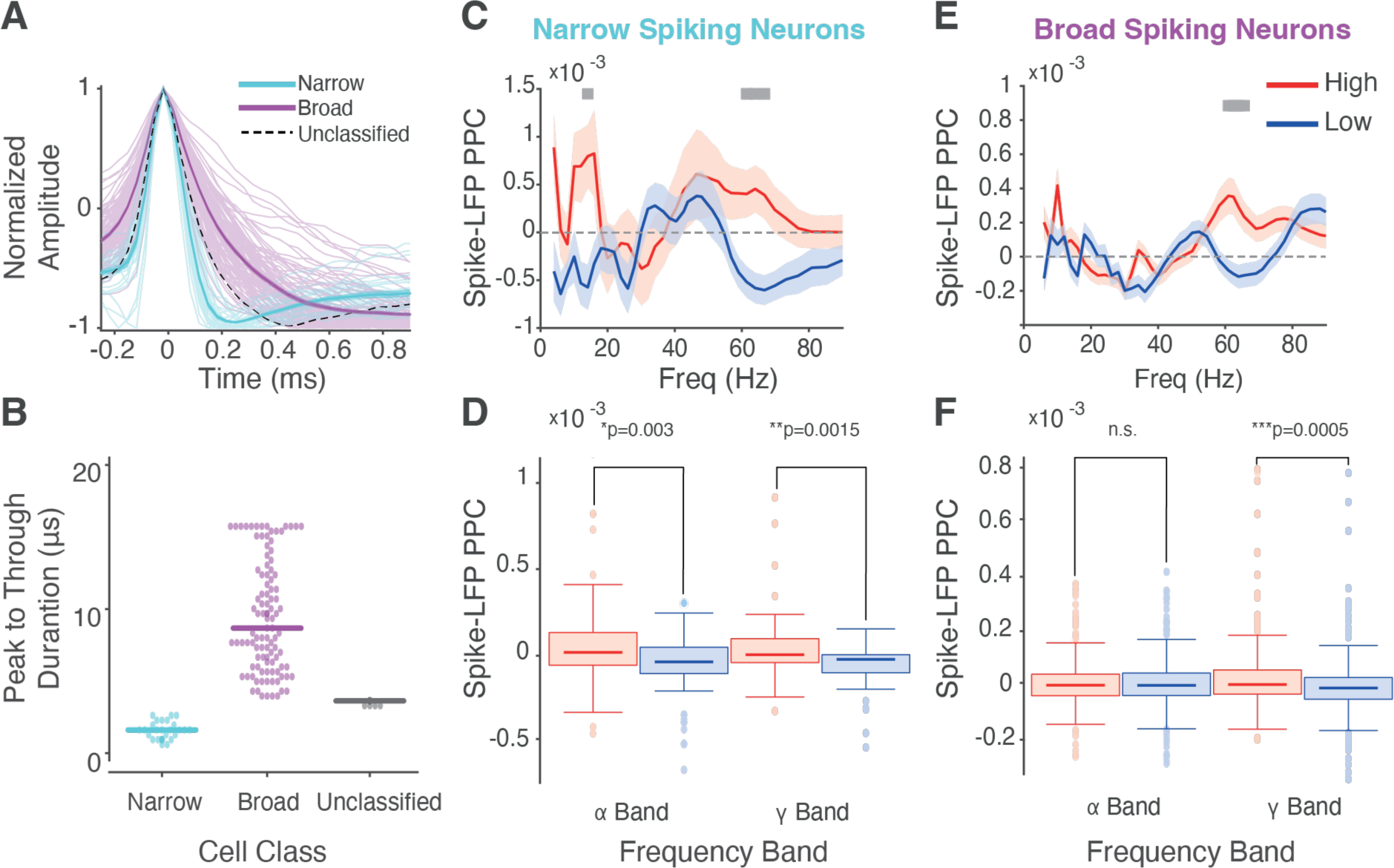
Spike-LFP phase coherence in primary visual cortex. Narrow- and broad-spiking cells increase their phase locking to alpha and gamma frequency-band during high-arousal quiescent states. **A)** Normalized spike waveform amplitude as a function of time (ms) for action potentials of neurons sorted by peak-to-trough-duration. Narrow (cyan), broad (violet), and unclassified (dashed line) spiking cells. Thick lines and ribbon area correspond to the average ± sem, respectively. **B)** Beeswarm plot of the neuronal cell types, with the horizontal line representing the median of each group. The observed distribution of the dots was bimodal (p=0.03, dip=0.04, Hartigan’s dip test). **C)** Pairwise phase consistency (PPC) spectrum of narrow-spiking cells sorted across high- (red) and low- QW (blue) arousal conditions. Average across V1 channels and narrow-spiking neurons ± sem (ribbon). Gray bars denote p < 0.05 corrected for multiple comparisons across frequencies, nonparametric randomization test. **D)** Box plot of PPC differences of narrow-spiking cells per band (alpha and gamma, based on the differences observed in *(C)* as a function of the high- and low arousal conditions (significance threshold: P < 0.05, Wilcoxon’s rank-sum test). **E)** Same as *(C)* but for broad-spiking cells. **F)** Same as *(D)* but for broad-spiking cells.

Next, we quantified the phase-locking consistency of the spikes from these two populations to the underlying LFP rhythm using the pairwise phase consistency (PPC) index across frequencies (Vinck et al. 2012). We observed that narrow and broad-spiking cells significantly increased their phase locking to different brain rhythms during high arousal. Narrow-spiking cells are simultaneously locked to an alpha (11 to 15 Hz) and a gamma (60-70 Hz) frequency band (Figure 7C, 11 to 15 Hz and 60 to 70 Hz, p<0.05 permutation test). In contrast, broad-spiking cells are only phase-locked to the 60-70 Hz gamma frequency band (Figure 7E, 60 to 70 Hz, p<0.05 permutation test). Figure 7D and 7F show the PPC average across the population of cells to these bands (Figure 7D, Exact Mann-Whitney U test: narrow-spiking population: alpha frequency band: p=0.003, U=2989; gamma frequency band: p=0.001, U=3030; broad-spiking population: alpha frequency band: p=0.95, U=53920; gamma frequency band: p=0.005, U=62179). These results suggest that high- and low-frequency brain rhythms distinctively influence the activity of visual cortex neurons during high arousal states, with low-frequency phase locking reflecting the engagement of narrow-spiking cells to an alpha rhythm that is likely associated with top-down modulation (Arnal et al. 2011; Kerkoerle et al. 2014; Bastos et al. 2015).

## Discussion

Our results indicate that the spontaneous arousal levels, gauged via pre-stimulus pupil variability, affects post-stimulus neuronal processing during quiescence in head-fixed ferrets passively observing visual stimuli. The pupil size of these ferrets showed spontaneous fluctuations over time (Figure 1), which seemed to correlate with different levels of arousal (Reimer et al. 2014; McGinley, David, et al. 2015; McGinley, Vinck, et al. 2015; Vinck et al. 2015; Einstein et al. 2017; Neske et al. 2019). When trials were grouped according to pre-stimulus pupil dilation, we observed that neurons in V1 increased their firing rate at their preferred orientation during high arousal states (Figure 2). Also, the LFP power amplitude shifted from a preeminence of low-frequency bands to higher frequencies. Stimulus onset shifted the peak of an alpha band (∼12 Hz) towards lower frequencies, but the magnitude of the peak shift depended on the arousal state (Figure 3). High arousal increased LFP-LFP phase relationships at lower frequencies within and between visual and parietal cortices (Figure 4). A computational model mimicking a laminar architecture receiving feedback from emulating arousal signals between V1 and PPC showed that this spectral signature is compatible with feedback from higher cortical areas targeting infragranular layers in PPc and supragranular layers in V1 (Figures 5 and 6). Finally, we observed that narrow and broad-spiking neurons phase-lock to different LFP rhythms when stimulated during high arousal. Broad-spiking neurons entrained to high-frequency oscillations (>60 Hz), whereas narrow-spiking neurons phase-locked to low (12-18 Hz) and high-frequency (50-80 Hz) rhythms (Figure 7C and 7E).

### Inactive to active states follow incremental changes in the variability and sensitivity of neuronal responses

Neuronal correlates of wakefulness have been studied in well-defined behavioral states of the sleep-wake cycle and during physical activity such as locomotion (Harris and Thiele 2011; McGinley, Vinck, et al. 2015; Olcese et al. 2016; Poulet and Crochet 2019). Active behavioral states produce a cortical desynchronization with a predominance of high-frequency oscillations, an increase in the variability of the neuronal activity, and an increase in the sensitivity of neurons in responding to sensory stimuli (Crochet et al. 2006; Gentet et al. 2010; Zagha et al. 2013; Reimer et al. 2014; McGinley, David, et al. 2015; Vinck et al. 2015, 2016; Poulet and Crochet 2019). Conversely, low-arousal states correlate with a general cortical synchronization, a prevalence of low-frequency rhythms, and decreased spike variability (Steriade et al. 1993, 2001; Amzica and Steriade 1997; Vyazovskiy et al. 2011; McCormick et al. 2014; Sanchez-Vives et al. 2017). QW shows similar but dampened features as those observed during locomotion or attention (Reimer et al. 2014; McGinley, Vinck, et al. 2015; Neske et al. 2019; Poulet and Crochet 2019) and recent studies have proposed that transitions from inactive to active states follow incremental changes in the variability and sensitivity of neuronal responses (McGinley, David, et al. 2015; Neske et al. 2019). This monotonic increase with arousal suggests the existence of a continuum between the range of neuronal responses and wakefulness states. Our results extend these findings, showing that pre-stimulus wakefulness affects neuronal responses to stimuli processing.

Our study focused on the trials with the smallest and largest pupil size during quiet wakefulness and determined that these two conditions unveil quantitative differences in neuronal sensitivity as a function of arousal in the ferret. High arousal QW increased stimulus selectivity in orientation-selective neurons in line with what was previously observed during locomotion compared to rest (Niell and Stryker 2010; Vinck et al. 2015) and QW compared to anesthesia (Ecker et al. 2014; Reimer et al. 2014). This sensitivity change of neurons in the primary visual cortex is not induced by a general increase in the spike rate (Figure 2A), a finding also observed in previous reports (Vinck et al. 2015; Fernandez et al. 2017; Poulet and Crochet 2019). Instead, we observed that these changes are associated with spike-count correlation differences among neurons (Figures 2B and 2C), together with changes in power amplitude and phase synchrony of the LFP signal.

The spike-count correlation accounts for the shared variability between pairs of two recorded neurons (Cohen and Kohn 2011). In the ferret primary visual cortex, we found that the shared variability across neurons transiently increases immediately after stimulus onset during high arousal trials. This is in contrast with previous studies showing a decrease in shared variability of spiking responses after stimulus onset (Renart et al. 2010; Renart and Machens 2014; Neske et al. 2019; Waschke et al. 2021). However, spike count correlations can also change as a function of wakefulness state, attention, or anesthesia (Reimer et al. 2014; Ruff and Cohen 2014; Snyder et al. 2014; Denfield et al. 2018), depending on the heterogeneity of the population under examination (Ecker et al. 2011; Arbab et al. 2018). Because neurons with similar orientation properties tend to have a higher degree of shared variability (Pachitariu et al. 2015), the transient high-arousal post-stimulus increase in spike-count correlations might reflect neurons with similar tuning properties. Moreover, we confirmed previous observations in V1 that describe a shift of the LFP power with increasing arousal levels (Figure 3; Crochet et al. 2006; Gentet et al. 2010; Bennett et al. 2013; Polack et al. 2013; Zagha et al. 2013; Reimer et al. 2014; Schneider et al. 2014; McGinley, David, et al. 2015; Vinck et al. 2015; Einstein et al. 2017; Fernandez et al. 2017; Stitt et al. 2018). We detected a peak shift of the alpha frequency band (∼11-15 Hz) towards lower frequencies in the primary visual cortex after stimulus onset (Figure 3). The magnitude of the shift depended on the wakefulness state of the animal (Figures 3C and 3D). The occipital alpha peak increases in frequency when subjects switch from passive viewing to an active cognitive task (Haegens et al. 2014). We found that while visual stimulation reduces the frequency peak of alpha power (Figure 3D), high arousal states show consistently higher alpha peaks than low arousal states. Previous studies focused on gamma oscillations have found that this peak variability might reflect rapid cyclic changes in synaptic excitation (and a proportional inhibitory counterbalance) within a cortical microcircuit (Atallah and Scanziani 2009; Spyropoulos et al. 2022). Our results show that this variability can also affect low frequency oscillatory components of the LFP.

Earlier studies have identified the activity of infragranular cortical layers and their thalamic modulation as the probable cortical source of alpha rhythms (Lopes da Silva and Leeuwen 1977; Lopes da Silva et al. 1980). This thalamic modulation sustains cortical connectivity during attentional states in primates and rodents (Saalmann et al. 2012; Schmitt et al. 2017). In ferrets, theta and alpha rhythms modulate the communication between the thalamus and area PPc depending on the wakefulness state of the animal (Stitt et al. 2018). We did not observe such parietal modulations, but unaltered responses in other cortical layers may have masked them. Future studies using brain-wide recording techniques might elucidate these questions.

### Intra and interareal phase synchronization fluctuates during quiescent wakefulness

Consistent local and long-range LFP-LFP phase relationships between areas associated with cognitive tasks have been described in several species (Bosman et al. 2014; Fries 2015; Vinck et al. 2016). A low frequency-band component dominates both intraparietal (Figure 4B) and parietal-visual LFP-LFP phase relationships (Figure 4C) in the ferret. This increase of low-frequency synchronization strength in periods of high arousal is in line with previous findings (Vyazovskiy et al. 2011; Olcese et al. 2016, 2018), but in addition to that we observed in V1 an increase in coherence (WPLI) in a narrow gamma frequency band. This narrow gamma-band (between 40 and 60 Hz) might represent the functional interaction between the primary visual cortex and the lateral geniculate nucleus, which also depends on the arousal state of the animal (Saleem et al. 2017; Schneider et al. 2020). Our results reveal frequency-specific influences between areas depending on the wakefulness state of the animal. In primate cortex, gamma influences have been reported to be systematically stronger in the feedforward direction whereas alpha and beta typically dominate in the feedback direction, and this organization is consistent with anatomical projections patterns (Bastos et al. 2015). We used a model that mimicked these frequency-specific interactions across areas. Our model suggests that frequency-specific interactions, and the effects of different arousal levels on such interactions, can be captured by straightforward firing-rate models with laminar connectivity (Mejias et al. 2016). The model predicts that top-down arousal signals target deep layers of PPc and superficial layers of V1. The modeling results are consistent with existing neuroanatomical connectivity patterns of macaque cortex (Markov et al. 2013, 2014): specific layers receiving top-down feedback projections depend on the hierarchical distance between source and target, with short/long hierarchical distances corresponding to deep/superficial layers, respectively. This cortical organization successfully explains the observed local and long-range functional connectivity interactions.

Besides modeling a probable mechanistic origin of the observed frequency-dependent interactions between visual and parietal areas, our computational model provides additional information regarding arousal signals. Assuming that the influence of high-arousal states on visual and parietal areas is conveyed via top-down signals from higher cortical areas, subcortical structures (e.g., thalamus), or neuromodulatory signals (Herrero et al. 2008), we have studied frequency-specific patterns emerging for different feedback configurations. Anatomical projections from higher to lower brain areas tend to be diffuse and unspecific (Markov et al. 2014). A computational model like the one we used allows for the exploration of specific projections that might play a role in the emergence of the observed dynamics. Of the four feedback types considered (Figure 6), feedback targeting deep layers of proximal areas (here: PPc) and superficial layers on distal areas (here: V1) can replicate the most notable features of the data, except for overestimating gamma oscillations in visual-parietal influences during high-arousal. Both connectivity patterns include some level of feedback to deep layers. Feedback connections targeting deep layers for both proximal and distal areas can correct this overestimation at the expense of deviating from the data in the low-frequency range for interactions within visual neurons (Figure 6B). Previous models have shown the importance of these feedback signals during top-down inhibition (Mejias et al. 2016), and realistic distributed representations during working memory (Mejias and Wang 2019), both being important concepts for predictive coding principles (Pennartz et al. 2019). Interestingly, these two feedback patterns are in line with the ones observed in feedback projections in macaque cortex (Markov et al. 2014) and mice (Harris et al. 2019), respectively. Our modeling results point to the hypothesis that cortical feedback in ferrets can include both scenarios, supporting a body of literature highlighting features that ferret cortex share with that of macaques and mice (Kaschube et al. 2010; Kaschube 2014).

### Neuronal entrainment to the low- and high-frequency LFP components during quiescent high-arousal reveals different modulatory effects over cortical microcircuits

Finally, we characterized the entrainment of primary visual cortex cell types, sorted according to their spike waveforms, to the observed cortical rhythms. We showed that putative excitatory and inhibitory neurons increase their entrainment to the LFP during high arousal states (Polack et al. 2013; Vinck et al. 2015). In addition, while broad-spiking cells (putative pyramidal neurons) phase-locked to high-frequency oscillations, narrow-spiking cells (putative interneurons) interacted with both high (60- 70 Hz) and low-frequency (10-15 Hz) oscillations. This phase-locking pattern is consistent with the role of narrow-spiking interneurons controlling cortical microcircuit computations (Isaacson and Scanziani 2011) and the generation of high-frequency oscillations (Cardin et al. 2009; Sohal et al. 2009; Veit et al. 2017). Recently, it has been observed that the action of specific parvalbumin-expressing interneurons might mediate cholinergic modulations within a cortical microcircuit (Garcia-Junco-Clemente et al. 2019). A subset of GABergic interneurons expressing the neuro-derived neurotrophic factor L1, receive cortico-cortical projections from neighboring microcircuits and control the gain of excitatory cells. These interneurons inhibit their excitatory cell dendrites and disinhibit their somata via parvalbumin activity (Malina et al. 2021). We hypothesize that the low-frequency entrainment of narrow-spiking cells observed in our data might reflect the mediation of several cortical and subcortical influences that arousal exerts over cortical microcircuits (Harris and Thiele 2011). Future studies must elucidate the role of the interneurons during quiet wakefulness.

In conclusion, our results show that the variability and sensitivity of cortical responses to a stimulus critically depend on the animal’s behavioral state before stimulus onset. Arousal states modulate intra- and interareal coherence among circuits, engaging local circuits with different LFP rhythms. Our analysis of high and low-arousal quiet wakefulness states supports the notion that behavioral states are associated with continuous changes in neuronal activity and dynamics across time, and provide further evidence that the variability observed during visual processing depends on the behavioral state prior to the stimulation.

## Acknowledgments

This work was supported by the FLAG-ERA Joint Transnational Call (JTC) - 2015 (project CANON) and JTC-2019 (project DOMINO) co-financed by the Netherlands Organization for Scientific Research (Nederlandse Organisatie voor Wetenschappelijk Onderzoek, NWO) to CAB, and the European Union’s Horizon 2020 Framework Program for Research and Innovation (grant numbers 720270—Human Brain Project SGA1 to CMAP, 785907—Human Brain Project SGA2 to CAB and CMAP, and 945539— Human Brain Project SGA3 to CMAP). The authors want to thank the members of the Technology Center of the University of Amsterdam for their help in building the electrophysiological setup.

## Conflict of interest

The authors report no conflict of interest.

Address correspondence to Conrado A. Bosman, Cognitive and Systems Neuroscience Group, Swammerdam Institute for Life Sciences, University of Amsterdam, Science Park 904, 1098XH Amsterdam, The Netherlands.

## Supplementary Figures

**Supplementary Figure 1.**
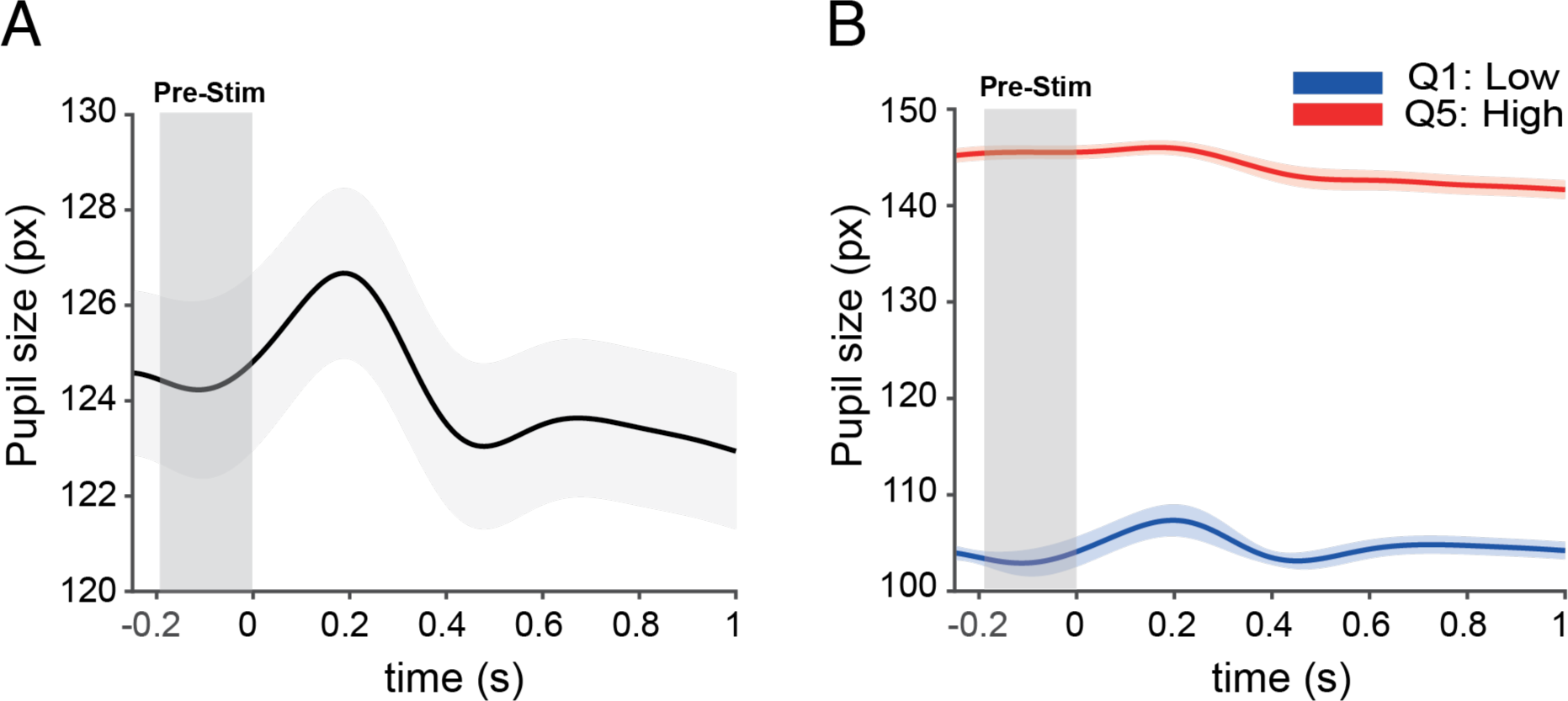
A) Pupil responses of one ferret (same as *Figure 1C*) but expressed as number of pixels (*px*). B) Pupil responses for the first quintile (Q1, low arousal) and the last quintile (Q5, high arousal) of one ferret (same as *Figure 1E*) expressed as number of pixels.

**Supplementary Figure 2.**
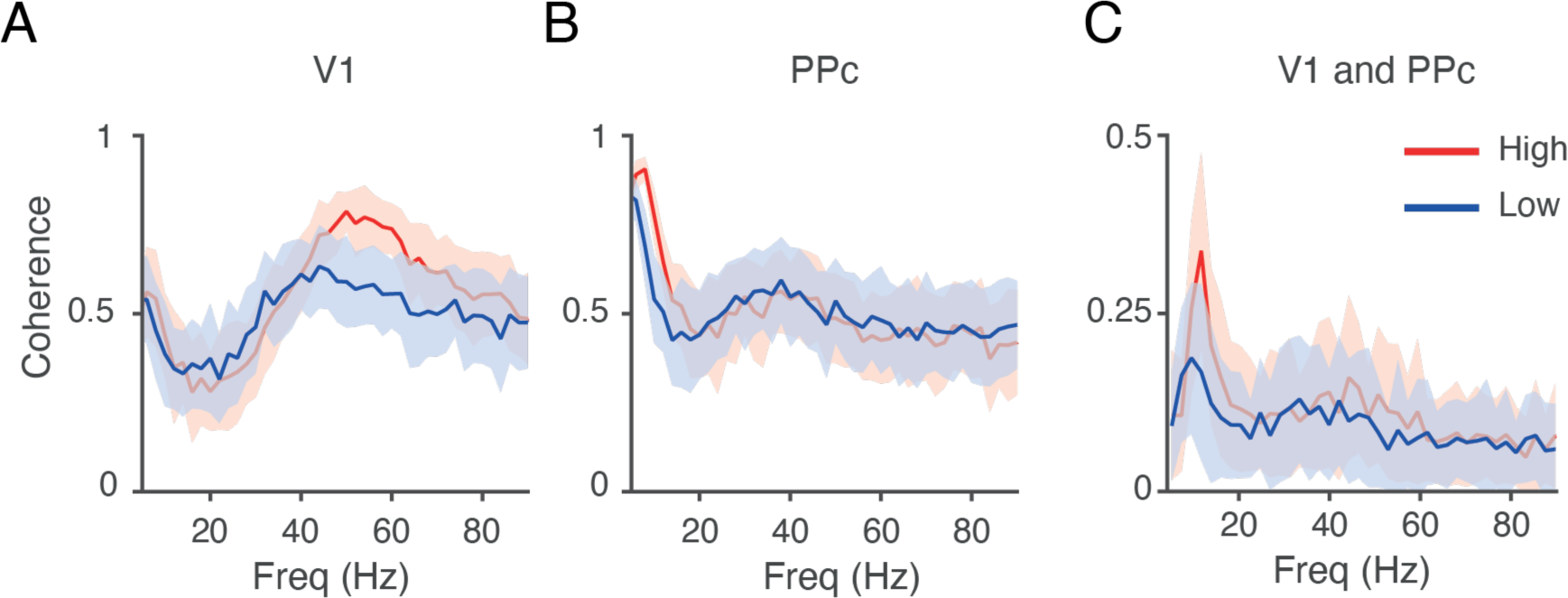
Same as *Figure 5*, but now using the coherence spectra as simulated by the model. **A to C.** Estimated coherence spectra from the model depicted in Figure 5A. **A)** Modeled coherence spectra of V1. **B)** Modeled coherence spectra of area PPc. **C)** Modeled coherence spectra between V1 and PPc. Mean ± standard deviation across different simulations. Color conventions as in Figure 5.

**Supplementary Figure 3.**
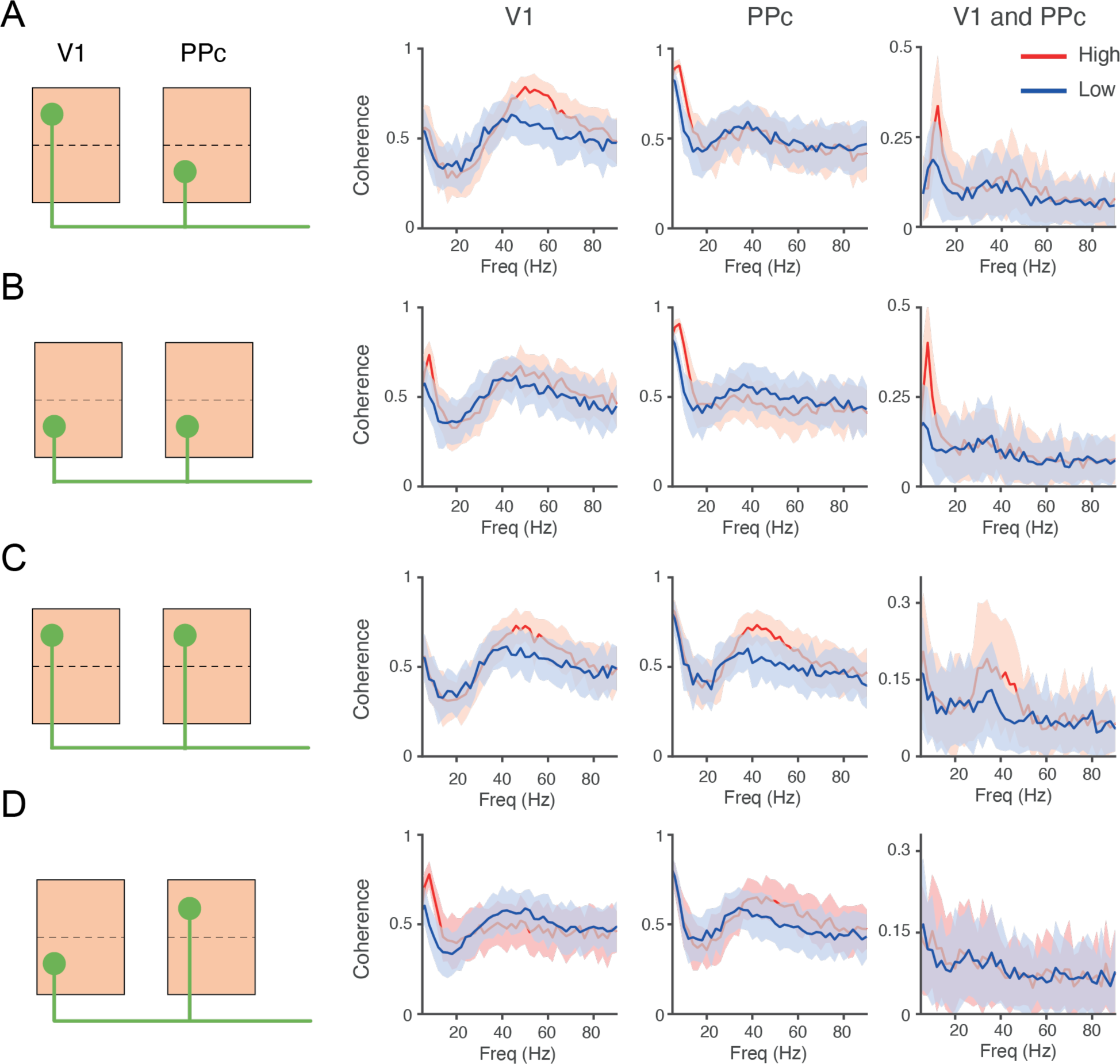
Inter- and intra-area coherence estimations vary according to the pattern of interlaminar projections in the computational model. **A to D)** *Left:* Variations in interlaminar projection patterns used to test the functional connectivity model. *Right:* Coherence spectra (using the WPLI index) between primary visual cortex and PPc (plotting conventions as *Figure 6*. Panel A is the same as *Supplementary Figure 2*).

## Notes

### Competing Interest Statement

The authors have declared no competing interest.

